# Divergent Retrograde Signaling Pathways Coordinate Longevity and Metformin Responses in Complex I and Complex IV Deficient *C. elegans*

**DOI:** 10.64898/2026.09.15.751667

**Authors:** Chutipong Chiamkunakorn, Juthakorn Poothong, Kongtana Trakarnsanga, Bart P. Braeckman, Wichit Suthammarak

## Abstract

Mild reductions in mitochondrial electron transport chain (ETC) capacity paradoxically extend organismal lifespan, a conserved phenomenon termed mitohormesis. While the mitochondrial unfolded protein response (UPR^mt^) and AMP-activated protein kinase (AMPK) are both established regulators of this process, whether distinct mitochondrial lesions converge on a single, unified survival mechanism remains unclear. By systematically dissecting the genetic architectures governing longevity in *Caenorhabditis elegans*, we demonstrate that targeted RNAi knockdown of Complex I (*nuo-6*) and Complex IV (*cco-1*) subunits activates fundamentally divergent downstream pathways. Both structural defects robustly induce the UPR^mt^, yet lifespan extension from Complex I impairment highly dependent on the UPR^mt^ master regulator ATFS-1 while bypassing the AMPK ortholog AAK-2. Conversely, Complex IV-mediated longevity operates independent of ATFS-1 but requires AAK-2-driven metabolic reprogramming. Pharmacological intervention with metformin—a Complex I inhibitor and AMPK activator—further exposed complex-specific vulnerabilities: metformin markedly suppressed the longevity phenotype of *nuo-6;aak-2* mutants, whereas in *cco-1;aak-2* animals it produced a trend toward lifespan extension that appeared independent of AAK-2. Together, these findings challenge the view of mitohormesis as a uniform response, revealing instead that cells engage specialized, molecularly tailored retrograde signaling networks to govern lifespan and respond to pharmacological interventions.

## Introduction

Aging is a fundamental physiological decline that acts as the primary risk factor for a spectrum of debilitating diseases, including neurodegeneration, cardiovascular disorders, and malignancies [1]. Extensive research into the molecular basis of longevity has identified several highly conserved nutrient-sensing and signal-transduction pathways—including insulin/insulin-like growth factor-1 (IGF-1) signaling, the mechanistic target of rapamycin (mTOR) pathway, AMP-activated protein kinase (AMPK), and NAD⁺-dependent sirtuins—together with mechanisms of translational and proteostatic homeostasis. These pathways frequently function as downstream effectors of systemic interventions such as caloric restriction, thereby modulating lifespan through coordinated regulation of cellular metabolism, stress resistance, and growth signaling [2–7]. Crucially, many of these lifespan-extending interventions converge upon mitochondrial function [8, 9]. Positioned at the critical nexus of cellular energy transduction, intermediary metabolism, and stress signaling, mitochondria serve as master regulators of both cellular vitality and organismal lifespan.

The intersection of mitochondrial bioenergetics and aging presents a profound biological paradox. While severe structural or biochemical disruptions to the mitochondrial electron transport chain (ETC) precipitate catastrophic bioenergetic crises and debilitating human pathologies, mild or tightly regulated reductions in respiratory capacity counterintuitively extend chronological lifespan. This evolutionary phenomenon, termed mitohormesis, demonstrates that sub-lethal curtailment of ETC performance delays senescent decline across diverse taxa, including yeast, flies, mice, and nematodes [9–14]. This observation remains deeply perplexing, as it suggests that compromised ATP synthesis and elevated reactive oxygen species (ROS) generation can delay, rather than accelerate, organismal deterioration [10, 15, 16].

To unravel this mitohormetic lifespan extension, the field has extensively utilized a distinct class of *Caenorhabditis elegans* respiratory mutants, particularly *nuo-6* [8, 9], which harbors a hypomorphic mutation in a core Complex I subunit, and *cco-1* [11, 15], which bears a disruption in an essential Complex IV structural subunit. These respiratory-deficient animals characteristically exhibit diminished metabolic rates, prolonged post-embryonic development, and significantly extended lifespans. Their longevity is largely attributed to the activation of retrograde signaling pathways, which transmit organellar stress to the nucleus to instigate cell-autonomous survival networks. A primary axis of this response is the mitochondrial unfolded protein response (UPR^mt^) [3], orchestrated by the bZIP transcription factor ATFS-1. During mitochondrial stress, ATFS-1 avoids local proteolysis, accumulates in the cytosol, and translocates to the nucleus to induce protective chaperones, such as HSP-6 [17–21]. Concurrently, mitochondrial respiratory disruptions elevate the cellular AMP/ATP ratio, stimulating AMP-activated protein kinase (AMPK, encoded by *aak-2* in *C. elegans*) [22], a metabolic master switch that shifts cellular resources toward catabolism and represses costly anabolic pathways to ensure survival during energetic deficits [16, 23–25]. Pharmacological strategies, most notably the biguanide metformin, frequently exploit these endogenous stress networks to promote healthy aging across model organisms, including *C. elegans* [26–28]. Metformin acts as a mild Complex I inhibitor [28–32], subsequently triggering downstream AMPK activation to augment longevity [26, 33]. Despite these mechanistic insights, a critical gap remains: historically, *nuo-6* and *cco-1* paradigms have been regarded as interchangeable models of mitochondrial longevity. However, because UPR^mt^ activation alone does not universally guarantee lifespan extension [34], this raises a fundamental research question: do distinct structural defects in the ETC converge on a unified survival mechanism, or do they trigger complex-specific retrograde signals?

In the present study, we hypothesized that mitochondrial dysfunction does not mediate lifespan extension through a single, generalized pathway. Rather, we posit that cells engage distinct, secondary signaling cascades tailored to the specific anatomical nature of the ETC defect. To test this, we systematically mapped the genetic architecture governing Complex I (*nuo-6*) and Complex IV (*cco-1*) longevity in *C. elegans*. We demonstrate that these defects activate fundamentally segregated pathways: Complex I impairment is highly dependent of the UPR^mt^ regulator ATFS-1, while Complex IV disruption bypasses ATFS-1 to rely heavily on AAK-2-driven metabolic reprogramming. Finally, by interrogating these distinct genetic backgrounds with the biguanide metformin, we expose specialized evolutionary vulnerabilities that dictate organismal survival and pharmacological response.

## Materials and Methods

### *C. elegans* Strains and Maintenance

The wild-type *Caenorhabditis elegans* strain N2 Bristol and the transgenic reporter strain SJ4100 (*zcIs13[hsp-6p*::GFP]) were utilized in this study. All nematode strains, alongside *Escherichia coli* (*E. coli*) HT115 (DE3) harboring the empty vector L4440 (EV) strains, were acquired from the Caenorhabditis Genetics Center (CGC, Minnesota, USA). Worms were cultured and maintained continuously at 20°C on standard Nematode Growth Medium (NGM) agar plates [35, 36]. All experimental protocols and animal handling procedures in this study were reviewed and approved by the Siriraj Animal Care and Use Committee (SIACUC), Faculty of Medicine Siriraj Hospital, Mahidol University, Bangkok, Thailand (Certificate of Approval No. 018/2562; Protocol No. SI-ACUP 003/2562).

### RNAi-Mediated Gene Knockdown

Gene silencing was executed via the standard bacterial feeding protocol [37]. Briefly, *E. coli* HT115 expressing double-stranded RNA (dsRNA), from Ahringer’s Library [38], targeting *cco-1* (F26E4.9), *nuo-6* (W01A8.4), *atfs-1* (ZC376.7), and *aak-2* (T01C8.1) were utilized. RNAi bacterial clones were cultured overnight at 37°C in Luria-Bertani (LB) broth supplemented with 100 µg/mL ampicillin. The bacterial cultures were subsequently seeded onto NGM plates containing 5 mM isopropyl 1-thio-β-D-galactopyranoside (IPTG) to induce dsRNA expression. For double RNAi knockdown experiments, the respective bacterial clones were mixed in equal volumes prior to seeding on the IPTG-supplemented NGM plates. Because this dual-RNAi feeding method inherently reduces the bacterial concentration of each targeted construct compared to single-RNAi treatments, knockdown efficacy was validated through functional epistasis, utilizing both downstream biochemical readouts (e.g., P-AAK-2 immunoblotting) and complete functional phenotypic reversion.

### Lifespan Assays

To initiate lifespan experiments, synchronized adult N2 worms were allowed to lay eggs for a period of 4 to 6 hours on NGM plates seeded with either empty vector (EV) control or the specific RNAi bacterial clones (*cco-1*, *nuo-6*, *atfs-1*, and *aak-2*). For pharmacological interventions, metformin was added to the NGM agar to a final concentration of 50 mM. All lifespan assays, including those involving metformin treatment, were conducted using live *E. coli* HT115 (DE3) harboring the empty vector L4440 (EV) to maintain continuous RNAi efficacy. To maintain cohort synchronization and prevent progeny contamination, worms were transferred to fresh plates daily during their reproductive phase. Each experimental cohort comprised approximately 100 synchronized animals. Mortality was defined as the complete failure to respond to gentle mechanical stimulation with a platinum wire. Worms that exhibited bagging, or those that crawled off the agar and were lost, were censored and excluded from the final experimental analysis. The resulting survival data were subsequently compiled and analyzed utilizing the Online Application for Survival Analysis (OASIS) open-source software (https://sbi.postech.ac.kr/oasis/) [39].

### Fluorescence Microscopy

For *in vivo* imaging of the *hsp-6p*::GFP reporter, live nematodes were immobilized and mounted on 2% agarose pads. Green fluorescence images were captured using a Nikon Ti-S inverted fluorescence microscope equipped with an Intensilight Ri1 camera and NIS-Elements D software (Nikon, Tokyo, Japan). Representative live-worm fluorescence images were captured under identical exposure settings across all conditions. Fluorescence intensity was subsequently quantified to statistically evaluate *hsp-6p*::GFP reporter expression across the experimental groups.

### Protein Extraction and Western Blotting

Total cellular proteins were extracted by homogenizing worms in RIPA lysis buffer (Thermo Fisher Scientific, catalog no. 89900), which ensures efficient solubilization of cytoplasmic, membrane, and nuclear proteins. Protein concentrations were determined using the Lowry protein assay kit (Bio-Rad, catalog no. 5000114). Twenty micrograms of total protein per sample were separated via 12% SDS-PAGE and subsequently transferred onto PVDF membranes. To block non-specific binding, membranes were incubated in 5% bovine serum albumin (BSA) dissolved in TBST buffer (Tris-buffered saline containing 0.1% Tween 20). Immunoblotting was performed using primary antibodies against Phospho-AMPKα (Thr172) (Cell Signaling Technology, catalog no. 2535) and β-actin (Cell Signaling Technology, catalog no. 8457), followed by an anti-rabbit IgG secondary antibody (Cell Signaling Technology, catalog no. 7074). Protein bands were visualized using a chemiluminescence ECL substrate (Bio-Rad, catalog no. 170-5060) and subjected to densitometric quantification to assess P-AAK-2 expression levels across conditions.

### Statistical Analysis

For lifespan assays, *p*-values were calculated using pairwise log-rank (Mantel-Cox) tests between specific experimental cohorts and their respective controls. Because comparisons were planned a priori based on distinct mechanistic hypotheses, *p*-values are reported without post-hoc adjustments for multiple comparisons. For the quantitative analysis of GFP fluorescence and Western blots, statistical significance was determined using an unpaired *t*-test (for comparing two groups) or a one-way ANOVA with appropriate post-hoc tests for multiple comparisons. All quantitative data are expressed as mean ± standard deviation (SD). Statistical significance is denoted as follows: ns: *p* > 0.05, *: *p* < 0.05, **: *p* < 0.01, ***: *p* < 0.001, ****: *p* < 0.0001

### Graphical Abstract and Figure Preparation

The graphical abstract, designed to illustrate the divergent retrograde signaling pathways and metabolic reprogramming architectures described in this study, was created using BioRender (https://biorender.com).

### Use of Generative AI Tools

During the preparation of this manuscript, the authors utilized Gemini (Google) strictly for language polishing, sentence structuring, and English prose editing to enhance readability. The AI tool was not used to collect, analyze, generate, or alter any raw data, experimental designs, or scientific conclusions. The authors reviewed, revised, and verified all edited content and take full responsibility for the accuracy and integrity of the final published manuscript.

## Results

### Mitochondrial Complex I and Complex IV Disruption Extends Lifespan and Induces the Mitochondrial Unfolded Protein Response (UPR^mt^)

To evaluate the physiological consequences of complex-specific mitochondrial electron transport chain (ETC) disruptions, we examined the lifespan and cellular stress signaling of *C. elegans* following RNAi-mediated knockdown of *nuo-6* (Complex I) and *cco-1* (Complex IV). Targeted mitochondrial impairment extended organismal lifespan compared to empty vector (EV) controls. EV animals exhibited a median lifespan of 17 days, whereas *nuo-6* knockdown provided a statistically significant lifespan extension, raising the median lifespan to 19 days (*p* = 0.036) (Figure 1A, Table 1). The most pronounced effect occurred with *cco-1* RNAi, which robustly extended median lifespan to 24 days, maintaining nearly 100% viability through day 15 (Figure 1A, Table 1). The greater magnitude of lifespan extension observed in *cco-1* relative to *nuo-6* knockdown is consistent with foundational mitohormesis models [9, 15]. Because long-lived mitochondrial mutants frequently activate protective retrograde signaling, we evaluated UPR^mt^ induction using an *hsp-6p*::GFP transcriptional reporter. While EV controls displayed negligible baseline fluorescence, both *nuo-6* and *cco-1* deficient animals exhibited a statistically significant quantitative upregulation of GFP, demonstrating a strong correlation between ETC disruption, UPR^mt^ activation, and enhanced longevity (Figure 1B-C).

**Figure 1.**
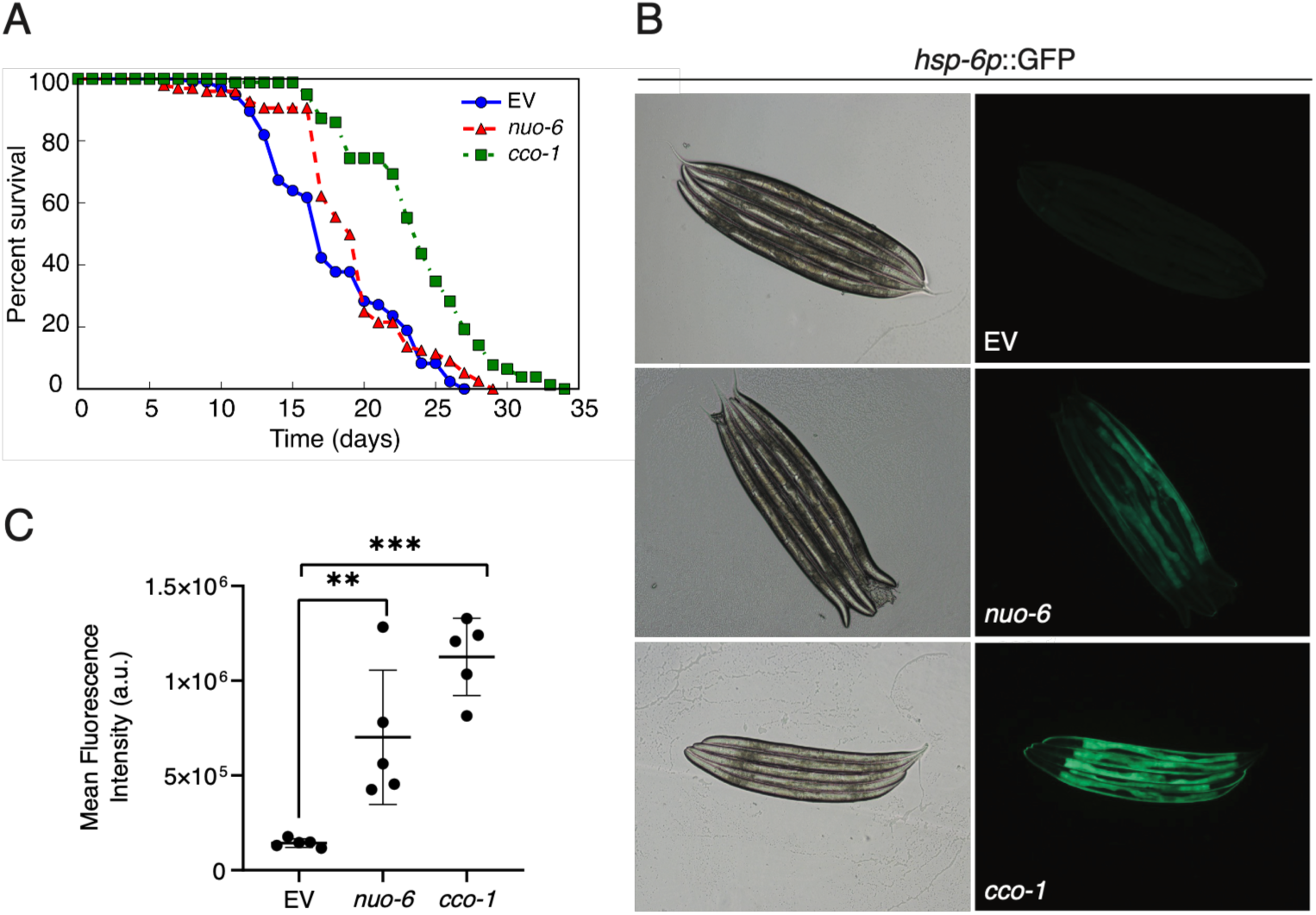
Disruptions of mitochondrial Complex I (*nuo-6*) and Complex IV (*cco-1*) extend lifespan and autonomously activate the mitochondrial unfolded protein response (UPR^mt^) in *C. elegans*. (A) Chronological lifespan curves of empty vector (EV) control, *nuo-6* RNAi, and *cco-1* RNAi animals. The *cco-1* and *nuo-6* knockdowns both display significant lifespan extension relative to EV controls (Table 1). (B) Representative brightfield (left) and fluorescence (right) micrographs of young adult *hsp-6p*::GFP reporter worms following EV, *nuo-6* , or *cco-1* RNAi treatment. Basal GFP fluorescence is minimal in EV controls but markedly induced upon *nuo-6* or *cco-1* depletion, indicating UPR^mt^ activation. Lifespan assays represent n = 95-100 animals per condition. Quantitative analysis of GFP fluorescence was evaluated using a one-way ANOVA with multiple comparisons. Data are presented as mean ± SD. (**: *p*<0.01, ***: *p*<0.001).

**Table 1.** Lifespan analysis of wild-type (EV) and genetic knockdown cohorts.

| Strain | n | Lifespan (days) |  |  | *p*-value | Comparison |
| --- | --- | --- | --- | --- | --- | --- |
|  |  | mean | median | max |  |  |
| EV | 100 | 17.89 | 17 | 27 |  |  |
| *atfs-1* | 100 | 17.98 | 17 | 30 | 0.835 | vs EV |
| *aak-2* | 100 | 16.55 | 17 | 26 | **0.023\*** | vs EV |
| *nuo-6* | 99 | 19.25 | 19 | 29 | **0.036\*** | vs EV |
| *nuo-6;atfs-1* | 100 | 14.71 | 15 | 25 | **<0.001\*** | vs *nuo-6* |
| *nuo-6;aak-2* | 95 | 20.37 | 21 | 29 | **0.014\*** | vs *nuo-6* |
| *cco-1* | 99 | 23.73 | 24 | 34 | **<0.001\*** | vs EV |
| *cco-1;atfs-1* | 100 | 23.49 | 24 | 31 | 0.982 | vs *cco-1* |
| *cco-1;aak-2* | 100 | 16.93 | 15 | 31 | **<0.001\*** | vs *cco-1* |

*p*-values were calculated using the log-rank test for pre-planned pairwise comparisons. The statistical significance level, denoted by \*, is considered when the *p*-value is less than 0.05.

### The Lifespan Extension of *nuo-6* and *cco-1* Deficient Worms Exhibits Differential Dependency on the Regulator ATFS-1

Given that ATFS-1 is the master nuclear regulator of the UPR^mt^, we investigated its role in orchestrating these stress responses and longevity phenotypes. Under baseline conditions, RNAi knockdown of *atfs-1* altered neither the survival kinetics (median 17 days) nor the basal *hsp-6p*::GFP fluorescence of wild-type worms (Figure 2A, 2D, 2G, Table 1). This baseline neutrality aligns with previous reports demonstrating that *atfs-1* depletion is phenotypically silent under basal conditions [18]. However, double RNAi-knockdown experiments revealed a profound mechanistic divergence between Complex I and Complex IV deficiencies. The lifespan extension conferred by *nuo-6* disruption was highly dependent on ATFS-1; upon *atfs-1* co-knockdown (*nuo-6;atfs-1*), the median lifespan of *nuo-6* animals dropped significantly from 19 to 15 days, accompanied by a near-complete ablation of *hsp-6p*::GFP fluorescence (Figure 2B, 2E, 2H, Table 1). Conversely, the robust longevity of *cco-1* animals proved independent of ATFS-1. Although *cco-1;atfs-1* mutants failed to induce the *hsp-6p*::GFP reporter, their survival kinetics mirrored those of *cco-1* single knockdowns, maintaining a prolonged median lifespan (Figure 2C, 2F, 2I, Table 1). Thus, ATFS-1 is broadly required for UPR^mt^chaperone transcription but is selectively essential for Complex I-mediated longevity.

**Figure 2.**
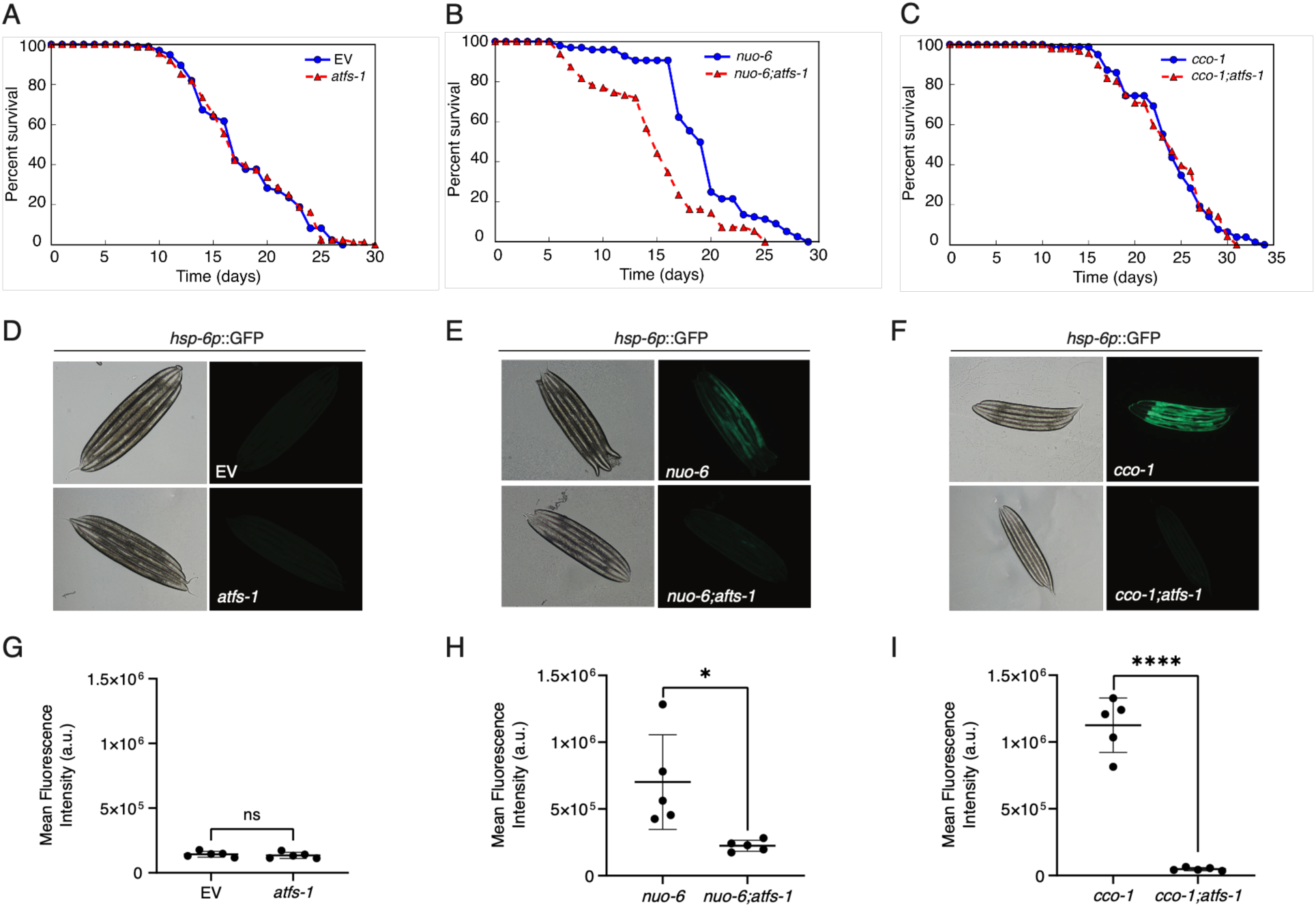
ATFS-1 is selectively required for Complex I (*nuo-6*)-mediated longevity but essential for UPR^mt^ activation in both *nuo-6* and *cco-1* knockdowns. (A) Chronological lifespan curves showing that *atfs-1* single knockdown does not alter baseline survival kinetics compared to EV control. (B) *atfs-1* co-knockdown significantly suppresses *nuo-6*-mediated lifespan extension. (C) Co-knockdown of *atfs-1* fails to suppress *cco-1*-mediated lifespan extension. (D) Representative fluorescence micrographs of *hsp-6p*::GFP animals showing reporter activation is unaltered in *atfs-1* single knockdown. (E) *atfs-1* co-knockdown profoundly suppressed *nuo-6*-induced *hsp-6p*::GFP reporter induction. (F) Co-knockdown of *atfs-1* eliminates *cco-1*-induced *hsp-6p*::GFP fluorescence. (G) Quantitative analysis of relative *hsp-6p*::GFP fluorescence intensity corresponding to the EV control and *atfs-1* single knockdown cohorts in panel D. (H) Quantitative analysis of relative *hsp-6p*::GFP fluorescence intensity demonstrating the profound suppression of the reporter signal in *nuo-6;atfs-1* double knockdowns compared to *nuo-6* single knockdowns. (I) Quantitative analysis of relative *hsp-6p*::GFP fluorescence intensity demonstrating the loss of reporter signal in *cco-1;atfs-1* double knockdowns compared to *cco-1* single knockdowns. Lifespan assays represent n = 95-100 animals per cohort; statistical significance was evaluated using the log-rank test (Table 1). Quantitative analysis of GFP fluorescence was evaluated using an unpaired *t*-test. Data are presented as mean ± SD. (ns: *p*>0.05 , *: *p*<0.05, ****: *p*<0.0001).

### The Energy Sensor AAK-2/AMPK Is Differentially Required for Complex IV-vs. Complex I-Mediated Lifespan Extension

To further delineate the metabolic networks coordinating these divergent longevity signals, we interrogated the cellular energy sensor AAK-2 (AMPK). Under standard conditions, *aak-2* knockdown moderately reduced overall mean survival and accelerated mortality following the 50% survival threshold compared to EV controls (Figure 3A, Table 1). This mild reduction in baseline resilience is consistent with prior reports demonstrating that AMPK deficiency impairs late-life metabolic homeostasis and accelerates mortality [22, 23]. When evaluating genetic dependencies, AAK-2 was dispensable for *nuo-6*-mediated longevity; in fact, *nuo-6;aak-2* double knockdowns exhibited a modest additional extension in median lifespan to 21 days compared to *nuo-6* single knockdowns (19 days, *p* = 0.014) (Figure 3B, Table 1). In stark contrast, the loss of *aak-2* profoundly suppressed the extended lifespan of *cco-1*-deficient animals. The *cco-1;aak-2* double knockdowns exhibited an accelerated onset of mortality, with their median lifespan decreasing from 24 days to 15 days (Figure 3C, Table 1). These data demonstrate that Complex I impairment promotes longevity via ATFS-1 while bypassing AAK-2, whereas Complex IV impairment acts independently of ATFS-1 but critically necessitates AAK-2-mediated metabolic reprogramming. To determine whether these genetic dependencies correspond to an altered cellular energy status, we evaluated AAK-2 phosphorylation (P-AAK-2)—a well-established biochemical indicator for an elevated cellular AMP/ATP ratio—via immunoblotting (Figure 3D). Densitometric quantification revealed that *cco-1* depletion significantly increased P-AAK-2 levels relative to EV control (*p* < 0.05), whereas *nuo-*6 depletion resulted in a non-significant trend toward reduced baseline AAK-2 levels *(p* > 0.05) (Figure 3E). These biochemical data strongly support our genetic epistasis findings, demonstrating that Complex IV impairment triggers an AAK-2-mediated energetic stress response to promote longevity, whereas Complex I-mediated longevity operates independently of this bioenergetic axis.

**Figure 3.**
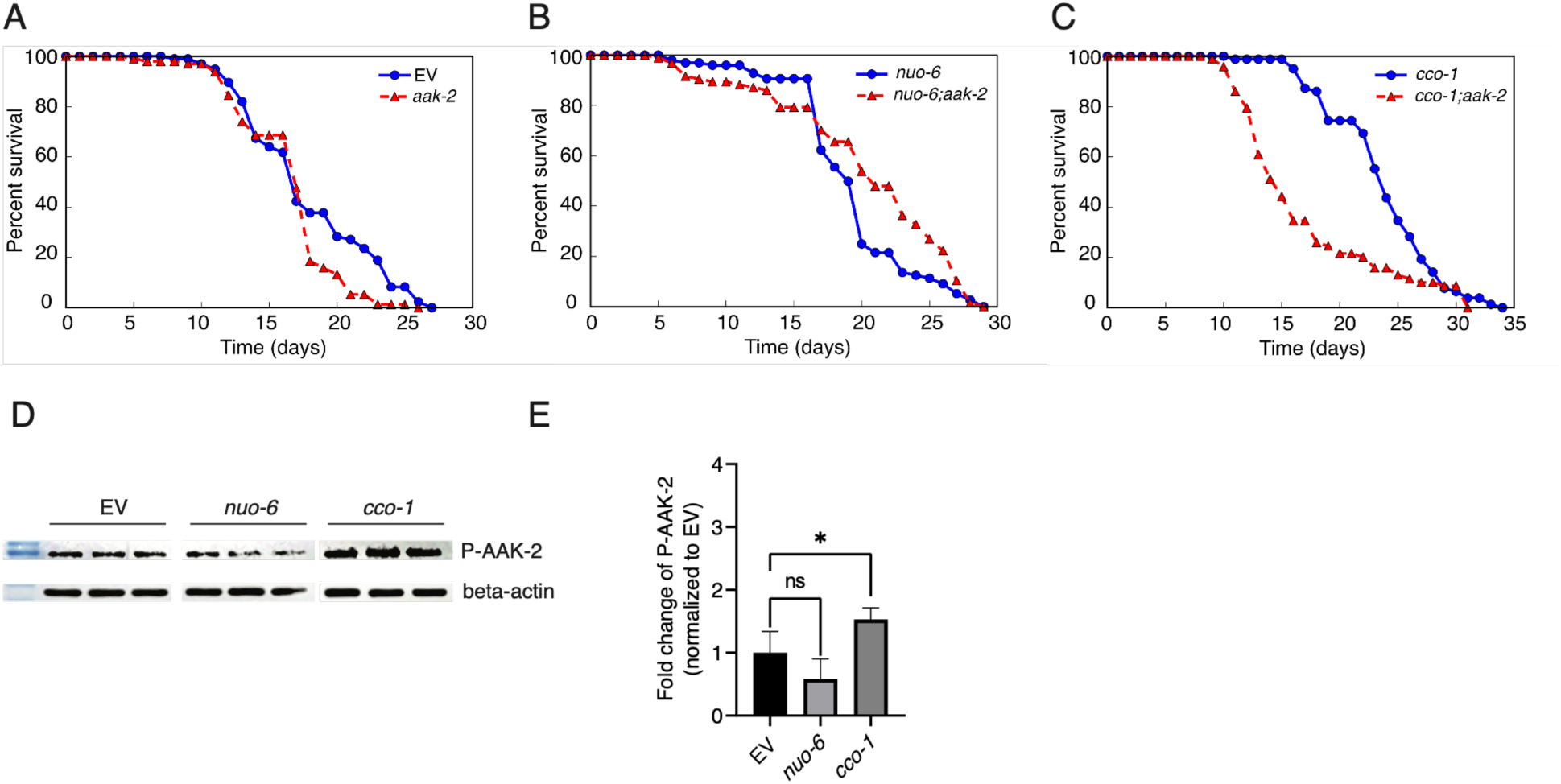
The energy sensor AAK-2/AMPK is selectively required for Complex IV (*cco-1*)- but dispensable for Complex I (*nuo-6*)-mediated longevity, where *aak-2* co-knockdown further extends survival. (A) Baseline chronological survival of EV control and *aak-2* RNAi animals, demonstrating that *aak-2* depletion causes a moderate reduction in baseline lifespan. (B) Lifespan analysis showing that *aak-2* co-knockdown does not compromise *nuo-6*-mediated lifespan extension (*nuo-6;aak-2*). (C) *aak-2* co-knockdown strongly suppressed *cco-1*-induced longevity (*cco-1;aak-2*), resulting in a severe truncation of median lifespan. (D) Representative Western blot of whole-worm lysates probed for phosphorylated AAK-2 (P-AAK-2) and β-actin as a loading control, illustrating AAK-2 activation status. (E) Quantitative densitometric analysis of relative P-AAK-2 band intensity from independent biological replicates (n= 3-4). Lifespan data represent n = 95-100 synchronized worms per cohort; *p*-values were calculated via the log-rank test (Table 1). Quantitative analysis of band intensity was evaluated using an unpaired *t*-test. Data are presented as mean ± SD. (ns: *p*>0.05, *: *p*<0.05).

### Metformin Activates AMPK and Elicits Divergent Lifespan Responses in Complex I and Complex IV Deficient Animals

To complement our genetic analyses, we utilized metformin—a biguanide and known Complex I inhibitor that activates AMPK—to probe pharmacological interactions with these pathways. Exposure of EV control worms to 50 mM metformin moderately prolonged lifespan (median 19 days) and increased AAK-2 phosphorylation (P-AAK-2), validating biochemical engagement of the energy-sensing cascade (Figure 4A-B, Table 2). To determine whether this pharmacological intervention provides additive benefits to genetic mitochondrial impairment, we evaluated metformin treatment in single knockdown models. While 50 mM metformin did not further enhance the lifespan of *nuo-6*-deficient animals (Figure 4C, Table 2), it induced a marked shift in the survival of *cco-1*-deficient worms compared to untreated controls, significantly extending the median lifespan to 30 days (Figure 4D, Table 2). We next assessed how this pharmacological stress interacts with genetic respiratory deficiencies in an *aak-2*-compromised background. In *nuo-6*-deficient worms, administering metformin to *nuo-6;aak-2* animals caused a reversal of lifespan extension; the median lifespan sharply truncated to 18 days (Figure 5A, Table 2). This indicates that layering a Complex I-inhibiting drug onto a genetic Complex I defect is highly detrimental in the absence of protective AAK-2 signaling. Intriguingly, a distinct phenotypic response emerged in *cco-1*-deficient worms. Treating *cco-1;aak-2* animals with metformin resulted in a trend toward lifespan extension independently of *aak-2*, shifting median survival to 20 days (*p* = 0.079) (Figure 5B, Table 2). Finally, immunoblot analyses validated these functional outcomes; exposure to 50 mM metformin significantly elevated P-AAK-2 fold-change in EV control animals (*p* < 0.01). In contrast, metformin exposure produced non-significant alterations in P-AAK-2 fold-change within ETC-deficient backgrounds, including a non-significant trend toward reduced P-AAK-2 in *nuo-6* single knockdowns (*p* > 0.05) and non-significant upward trends in *cco-1* (*p* > 0.05) and *cco-1;aak-2* cohorts (*p* > 0.05), while the *nuo-6;aak-2* cohort displayed no fold-change in P-AAK-2 levels (Figure 5C-F).

**Figure 4.**
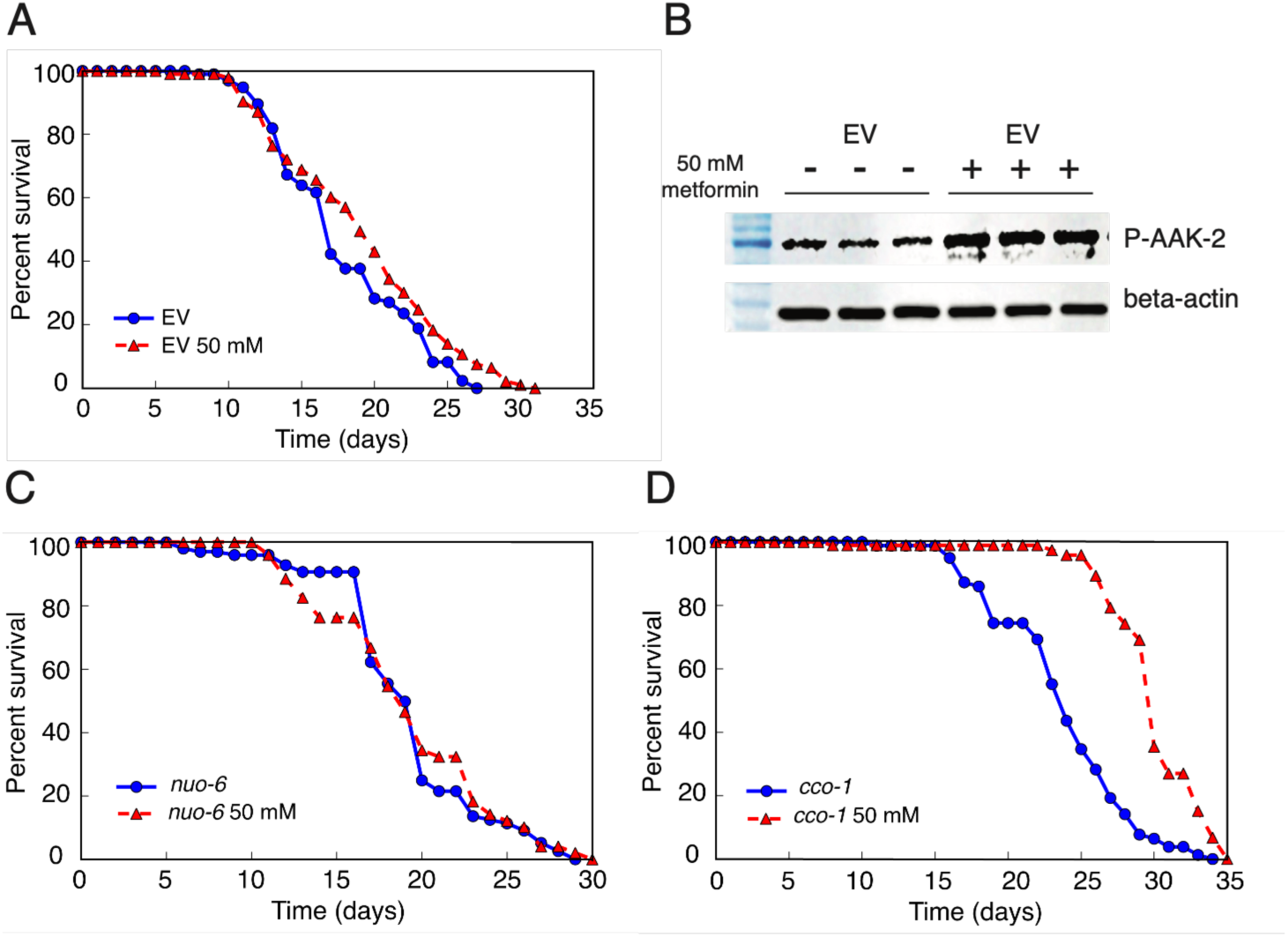
Metformin extends lifespan and stimulates AAK-2 (AMPK) phosphorylation in wild-type animals, and differentially affects longevity in Complex I- and IV-deficient worms. (A) Chronological lifespan curves of EV control worms treated with or without 50 mM metformin, showing a significant extension in median and maximum lifespan (*p* < 0.05, log-rank test; Table 2). (B) Representative Western blot of whole-worm lysates from EV control animals exposed to 0 mM (-) or 50 mM (+) metformin. Immunoblots were probed for phosphorylated AAK-2 (P-AAK-2) and β-actin as a loading control, confirming biochemical activation of AMPK signaling following treatment. (C) Chronological lifespan curves of *nuo-6* RNAi animals treated with 0 mM or 50 mM metformin, demonstrating that metformin treatment does not additionally extend lifespan in a Complex I-deficient background (*p* > 0.05, log-rank test; Table 2). (D) Chronological lifespan curves of *cco-1* RNAi animals treated with 0 mM or 50 mM metformin, illustrating significant lifespan extension upon metformin exposure in a Complex IV-deficient background (*p* < 0.001, log-rank test; Table 2).

**Figure 5.**
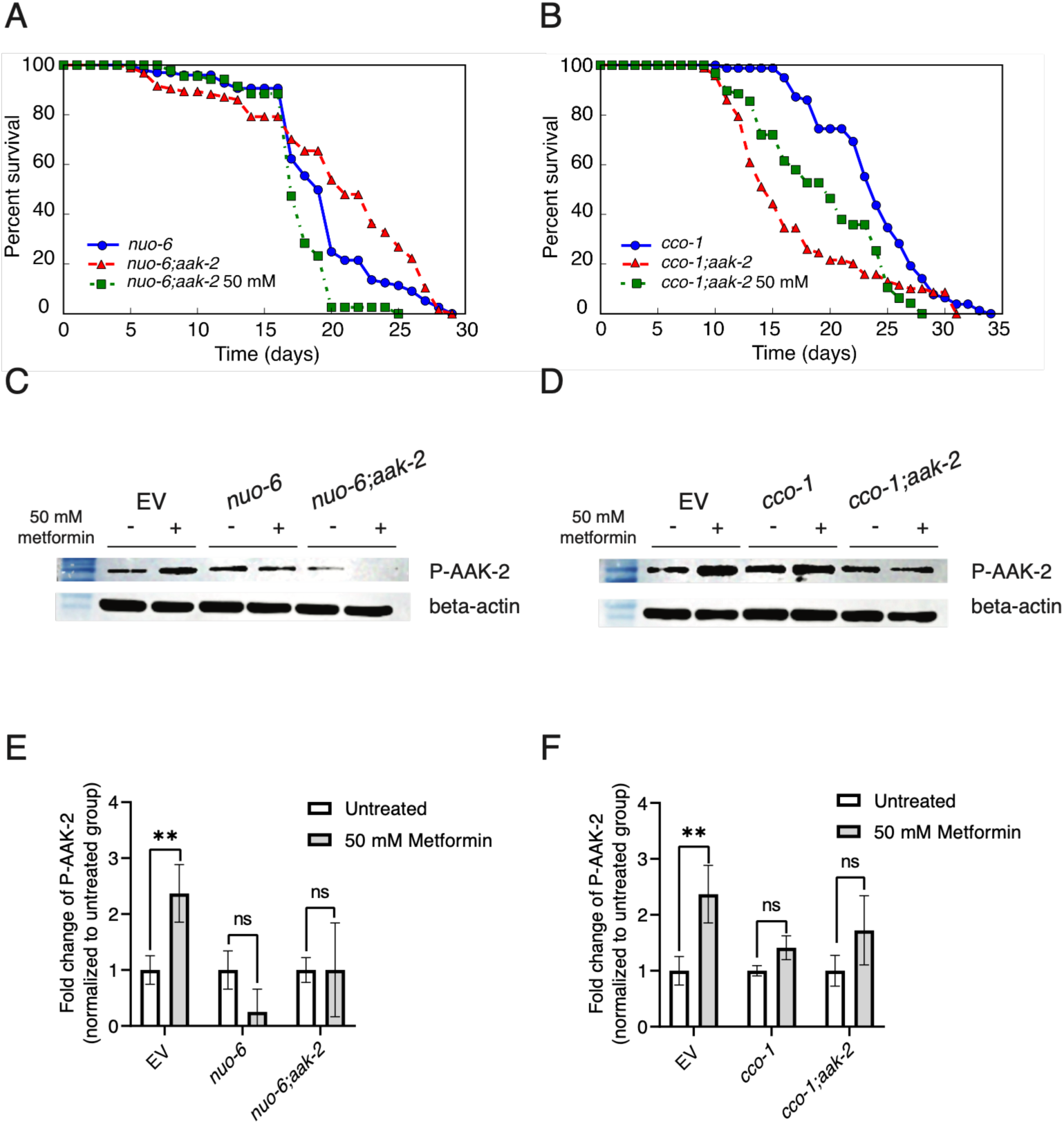
Metformin reverses longevity in *aak-2*-deficient Complex I knockdowns but extends lifespan in Complex IV mutants independently of *aak-2*. (A) Chronological lifespan curves showing that treating *nuo-6;aak-2* double-knockdown worms with 50 mM metformin ablates their extended lifespan phenotype (*p* < 0.001, log-rank test; Table 2). (B) Lifespan analysis showing that 50 mM metformin treatment shifts median survival of *cco-1;aak-2* double-knockdown animals from 15 to 20 days (*p* = 0.079, log-rank test; Table 2). (C) Western blot analysis of whole-worm lysates from EV control, *nuo-6*, and *nuo-6;aak-2* cohorts treated with (+) or without (-) 50 mM metformin. (D) Western blot analysis of whole-worm lysates from *cco-1* and *cco-1;aak-2* cohorts treated with (+) or without (-) 50 mM metformin. Blots probed for P-AAK-2 and β-actin show partial ablation of the P-AAK-2 signal upon *aak-2* knockdown, demonstrating that metformin fails to fully activate downstream energy-sensing cascades in these backgrounds. (E, F) Quantitative densitometric analysis of relative P-AAK-2 levels for the cohorts shown in panels C and D from independent biological replicates (n=3-4). To isolate the effect of metformin, the band intensity for each metformin-treated cohort is expressed as a fold-change relative to its own untreated baseline (set to 1.0). Statistical significance between two groups was evaluated using an unpaired *t*-test. Data are presented as mean ± SD. (ns: *p* > 0.05, **: *p* < 0.01).

**Table 2.** Lifespan analysis of wild-type (EV) and genetic knockdown cohorts treated with or without metformin.

| Strain | n | Metformin | Lifespan (days) |  |  | *p*-value | Comparison |
| --- | --- | --- | --- | --- | --- | --- | --- |
|  |  |  | mean | median | max |  |  |
| EV | 100 | 0 mM | 17.89 | 17 | 27 |  |  |
| EV | 100 | 50 mM | 19.14 | 19 | 31 | **0.046\*** | vs untreated EV |
| *nuo-6* | 99 | 0 mM | 19.25 | 19 | 29 | **0.036\*** | vs untreated EV |
| *nuo-6* | 100 | 50 mM | 19.27 | 19 | 30 | **0.768** | vs untreated *nuo-6* |
| *nuo-6;aak-2* | 95 | 0 mM | 20.37 | 21 | 29 | **0.014\*** | vs untreated *nuo-6* |
| *nuo-6;aak-2* | 100 | 50 mM | 17.46 | 18 | 25 | **<0.001\*** | vs untreated *nuo-6;aak-2* |
| *cco-1* | 99 | 0 mM | 23.73 | 24 | 34 | **<0.001\*** | vs untreated EV |
| *cco-1* | 100 | 50 mM | 29.97 | 30 | 35 | **<0.001\*** | vs untreated *cco-1* |
| *cco-1;aak-2* | 100 | 0 mM | 16.93 | 15 | 31 | **<0.001\*** | vs untreated *cco-1* |
| *cco-1;aak-2* | 100 | 50 mM | 19.32 | 20 | 28 | 0.079 | vs untreated *cco-1;aak-2* |

*p*-values were calculated using the log-rank test for pre-planned pairwise comparisons. The statistical significance level, denoted by \*, is considered when the *p*-value is less than 0.05.

## Discussion

The evolutionary paradox of mitohormesis posits that sublethal mitochondrial electron transport chain (ETC) disruptions can counterintuitively extend lifespan [40–43]. Historically, the aging field has frequently treated different mitochondrial mutants as interchangeable models [8], often positioning the mitochondrial unfolded protein response (UPR^mt^) as a universal driver of ETC-mediated longevity [3]. Our findings challenge this monolithic view. While targeted RNAi knockdown of both *nuo-6* (Complex I) and *cco-1* (Complex IV) robustly triggered the UPR^mt^, the downstream survival benefits fundamentally diverged. We found that the UPR^mt^ master regulator ATFS-1 is critically required for the longevity of *nuo-6* animals, whereas *cco-1*-mediated lifespan extension is entirely ATFS-1 independent. This critical distinction aligns with emerging evidence that UPR^mt^ activation, while a hallmark of mitochondrial stress, does not universally guarantee lifespan extension [34, 44]. Mechanistically, this divergence suggests that Complex I defects may induce a specific threshold of mitochondrial protein import stress that prevents ATFS-1 degradation, obligating its nuclear translocation to drive proteostatic survival networks. Conversely, structural perturbations at Complex IV likely trigger alternative retrograde signals that bypass the need of this proteostatic axis for longevity. This functional dichotomy aligns perfectly with and synthesizes independent observations previously published. For instance, while ATFS-1 was shown to mediate stress resistance and longevity in *nuo-6* mutants [45], other studies noted that *cco-1* RNAi retains its longevity benefits even when *atfs-1* is deleted [34]. Our data unifies these previously disjointed findings into a cohesive model: survival mechanisms are dictated by the specific site of the ETC lesion.

Having established that Complex IV-mediated longevity bypasses ATFS-1, we identified the AMP-activated protein kinase (AMPK) ortholog, AAK-2, as its essential mechanistic driver [46–48]. While AAK-2 is a well-established guardian of cellular energy homeostasis and metabolic reprogramming during energetic stress [25, 47], our study reveals its distinct necessity for Complex IV, but not Complex I, mutant survival. We hypothesize that this dichotomy is rooted in discrete bioenergetic crises dictated by the anatomical site of the ETC block. As the terminal electron acceptor, Complex IV dysfunction may cause a more severe bottleneck in oxidative phosphorylation. Our indirect assessment via P-AAK-2 densitometry strongly supports a proposed mechanism where this severe bottleneck elevates the cellular energetic deficit, making survival highly reliant on AAK-2-mediated shifts toward catabolism [49–52]. This aligns with established paradigms where profound metabolic remodeling is obligatory for survival during severe energetic crises at terminal OXPHOS sites [3, 47]. In contrast, Complex I defect might permit compensatory electron entry via alternative routes (such as Complex II) [53], thereby mitigating the acute energetic collapse and bypassing the obligate need for the AAK-2 metabolic switch.

Recent unbiased RNA-seq profiles of *C. elegans* mitochondrial mutants reveal that ETC disruptions do not trigger an identical transcriptional response [54]. For instance, transcriptomic profiling of *cco-1* knockdown animals reveals extensive shifts in chromatin remodeling, innate immunity, and metabolic restructuring that extend far beyond canonical chaperone induction [55]. In contrast, the transcriptome of *nuo-6* mutants is distinctly anchored by high-confidence ATFS-1 target genes involved in protein folding and targeted stress resistance [56]. These independent *in silico* datasets are broadly consistent with our *in vivo* genetic findings: *nuo-6* relies heavily on the ATFS-1 proteostatic axis, whereas the profound metabolic restructuring required for *cco-1* survival necessitates broader catabolic regulators like AAK-2. Because these transcriptomic datasets derive from independent studies rather than the same genetic backgrounds used here, we regard them as corroborating context rather than direct validation of our epistasis models.

The specialization of these retrograde networks is further illustrated by our pharmacological interventions with metformin, a mild Complex I inhibitor and downstream AMPK activator [28]. In an energy-sensing–compromised background (*nuo-6;aak-2* mutants), metformin ablated the extended lifespan phenotype. Metformin is known to promote longevity and metabolic resilience in *C. elegans* through LKB1/AMPK-dependent signaling pathways [26]. However, combining a pharmacological Complex I inhibitor onto a genetic Complex I defect likely triggers a localized ‘double-hit’—a compounded, intolerable bioenergetic crisis at a single mitochondrial site. Lacking the protective AAK-2 network to curtail energy expenditure, this localized stress overwhelms the organism, and the severity of this collapse likely hinders any potential rescue by secondary pleiotropic networks. Intriguingly, a fundamentally distinct phenotypic response emerged in Complex IV-deficient worms. While our genetic models demonstrate that baseline *cco-1*-mediated longevity strongly requires AAK-2, metformin treatment produced a trend toward extended lifespan in *cco-1;aak-2* double knockdowns. We propose that because metformin targets Complex I, the spatially distinct structural defect across the electron transport chain in Complex IV mutants diffuses this acute ‘double-hit’ toxicity. Consequently, metformin can exert its well-documented pleiotropic effects to artificially bypass the canonical AAK-2 metabolic requirement. Specifically, it may optimize alternative electron flow or engage independent, off-target mitohormetic stress responses—such as PRDX-2 activation or SKN-1/Nrf2 oxidative stress resistance [26, 28], or dietary restriction-like signaling—thereby conferring survival benefits independently of canonical AAK-2 energy sensing. Importantly, our assays utilized live *E. coli* HT115 to facilitate continuous RNAi. Metformin has been robustly demonstrated to alter bacterial metabolism—specifically the microbial folate and methionine cycles—and can induce a bacteriostatic dietary restriction state that extends *C. elegans* lifespan [57]. Therefore, the unexpected trend toward lifespan extension in metformin-treated *cco-1;aak-2* animals may be partially driven by these drug-microbe-host interactions, providing an exogenous survival signal that bypasses the worm’s canonical AAK-2 energetic sensor.

While our study clearly delineates divergent signaling requirements, we acknowledge certain methodological limitations. First, our genetic interaction studies relied on dual-RNAi feeding, which inherently reduces the bacterial concentration of each targeted construct compared to single-RNAi treatments. However, we address this potential caveat by anchoring our conclusions strictly on robust functional epistasis. The complete, quantitative ablation of the *hsp-6p*::GFP reporter fluorescence, coupled with profound shifts in survival kinetics and P-AAK-2 immunoblot signals across the double-knockdown cohorts, represents profound phenotypic reversion. These robust responses strongly argue against incomplete construct delivery driving the observed mechanistic dichotomies, confirming that knockdown efficacy remained high. Nonetheless, future studies utilizing stable genetic null mutants (e.g., *atfs-1* or *aak-2* deletion strains) will be valuable to further validate these epistatic relationships. Second, while our robust lifespan phenotypes and functional epistasis— supported by P-AAK-2 phosphorylation readouts—provide compelling indirect evidence of energetic stress, future investigations should incorporate direct biochemical profiling. Quantifying absolute cellular AMP/ATP ratios, individual ETC complex enzymatic activities, and ROS dynamics will be critical to definitively validate the distinct bioenergetic crises and mechanistic hypotheses generated by our genetic models.

## Conclusion

Mitochondrial longevity is not governed by a single, unified pathway but by anatomically tailored retrograde signaling networks. Complex I impairment relies on the ATFS-1-mediated UPR^mt^ activation, while Complex IV disruption requires AAK-2-driven metabolic reprogramming. Consequently, pharmacological interventions such as metformin must be precisely contextualized against preexisting mitochondrial lesions to avoid negating the organism’s innate survival benefits, underscoring the need for pathway-specific therapeutic targeting in aging and metabolic disease.

## Supporting information

Lifespan and Western Blot

## Acknowledgements

This work is supported by National Research Council of Thailand (NRCT) and Mahidol University (NRCT5-RSA63015-20). Worm strains were provided by the CGC, which is funded by NIH Office of Research Infrastructure Programs (P40 OD01044)

## Author Contributions

**Chutipong Chiamkunakorn:** Methodology, Formal analysis, Investigation, Writing - Original Draft. **Juthakorn Poothong:** Methodology, Investigation. **Kongtana Trakarnsanga:** Resources, Methodology. **Bart P. Braeckman:** Conceptualization, Writing - Review & Editing. **Wichit Suthammarak:** Conceptualization, Supervision, Funding acquisition, Project administration, Writing - Review & Editing.

## Data Availability Statement

The data that support the findings of this study are available in the supplementary material of this article.

## Abbreviations

AAK-2: AMP-activated protein kinase alpha subunit (C. elegans ortholog)
AMPK: AMP-activated protein kinase
ATFS-1: Activating Transcription Factor for Stress response-1
ATP: Adenosine triphosphate
CGC: Caenorhabditis Genetics Center
dsRNA: Double-stranded RNA
ETC: Electron transport chain
EV: Empty vector
GFP: Green fluorescent protein
IGF-1: Insulin/insulin-like growth factor-1
IPTG: Isopropyl 1-thio-β-D-galactopyranoside
LB: Luria-Bertani
NGM: Nematode Growth Medium
OXPHOS: Oxidative phosphorylation
ROS: Reactive oxygen species
T2DM: Type 2 diabetes mellitus
UPRmt: Mitochondrial unfolded protein response

---

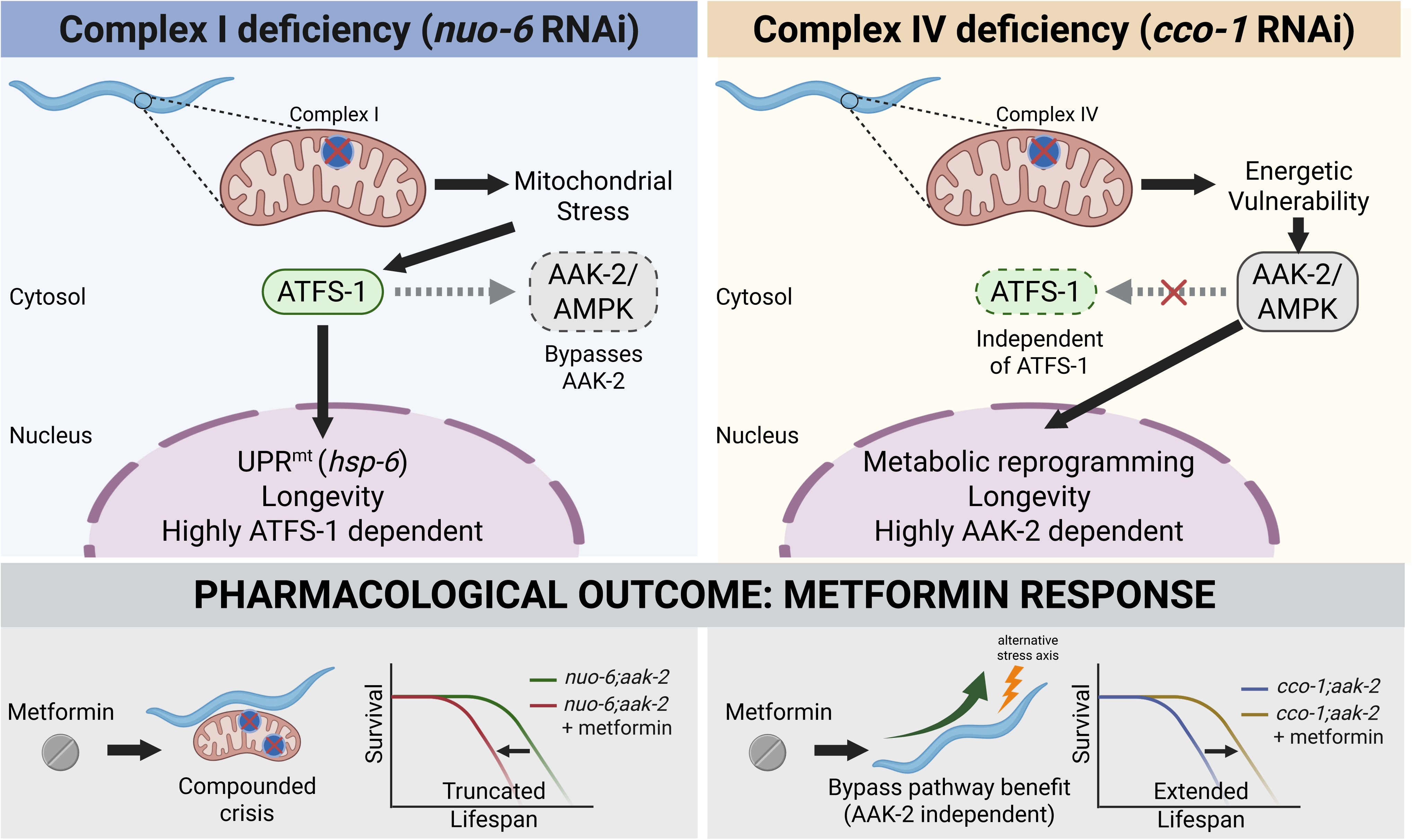

