## Supplementary material for "Divergent Retrograde Signaling Pathways Coordinate Longevity and Metformin Responses in Complex I and Complex IV Deficient *C. elegans*": Lifespan and Western Blot: Blot for Fig 5.pptx

### Slide 1
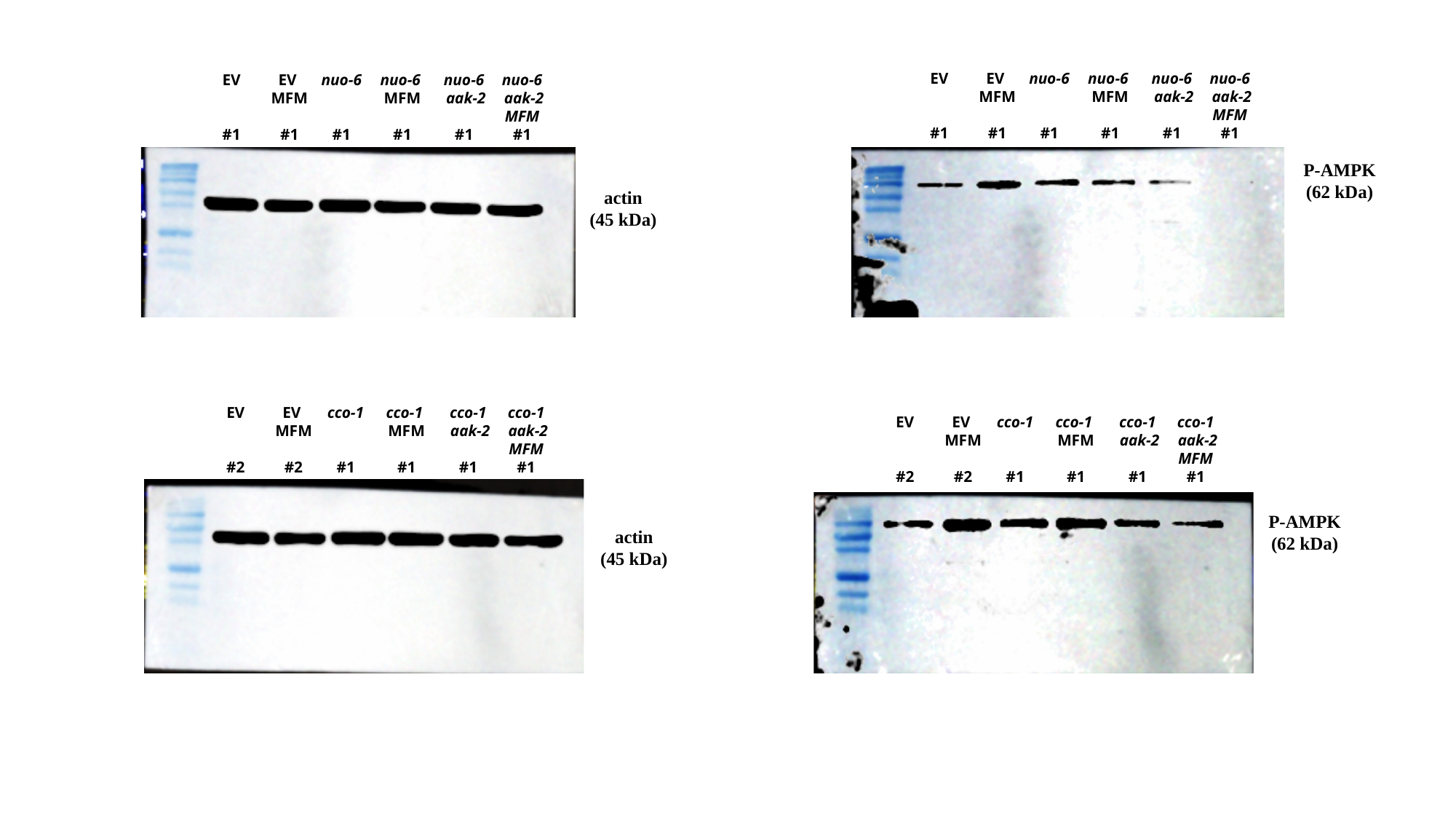

EV
#1
EV
MFM
#1
nuo-6
#1
nuo-6
MFM
#1
nuo-6
 aak-2
#1
nuo-6
 aak-2
MFM
#1
EV
#1
EV
MFM
#1
nuo-6
#1
nuo-6
MFM
#1
nuo-6
 aak-2
#1
nuo-6
 aak-2
MFM
#1
P-AMPK
(62 kDa)
EV
#2
EV
MFM
#2
cco-1
#1
cco-1
MFM
#1
cco-1
 aak-2
#1
cco-1
 aak-2
MFM
#1
EV
#2
EV
MFM
#2
cco-1
#1
cco-1
MFM
#1
cco-1
 aak-2
#1
cco-1
 aak-2
MFM
#1
P-AMPK
(62 kDa)

### Slide 2
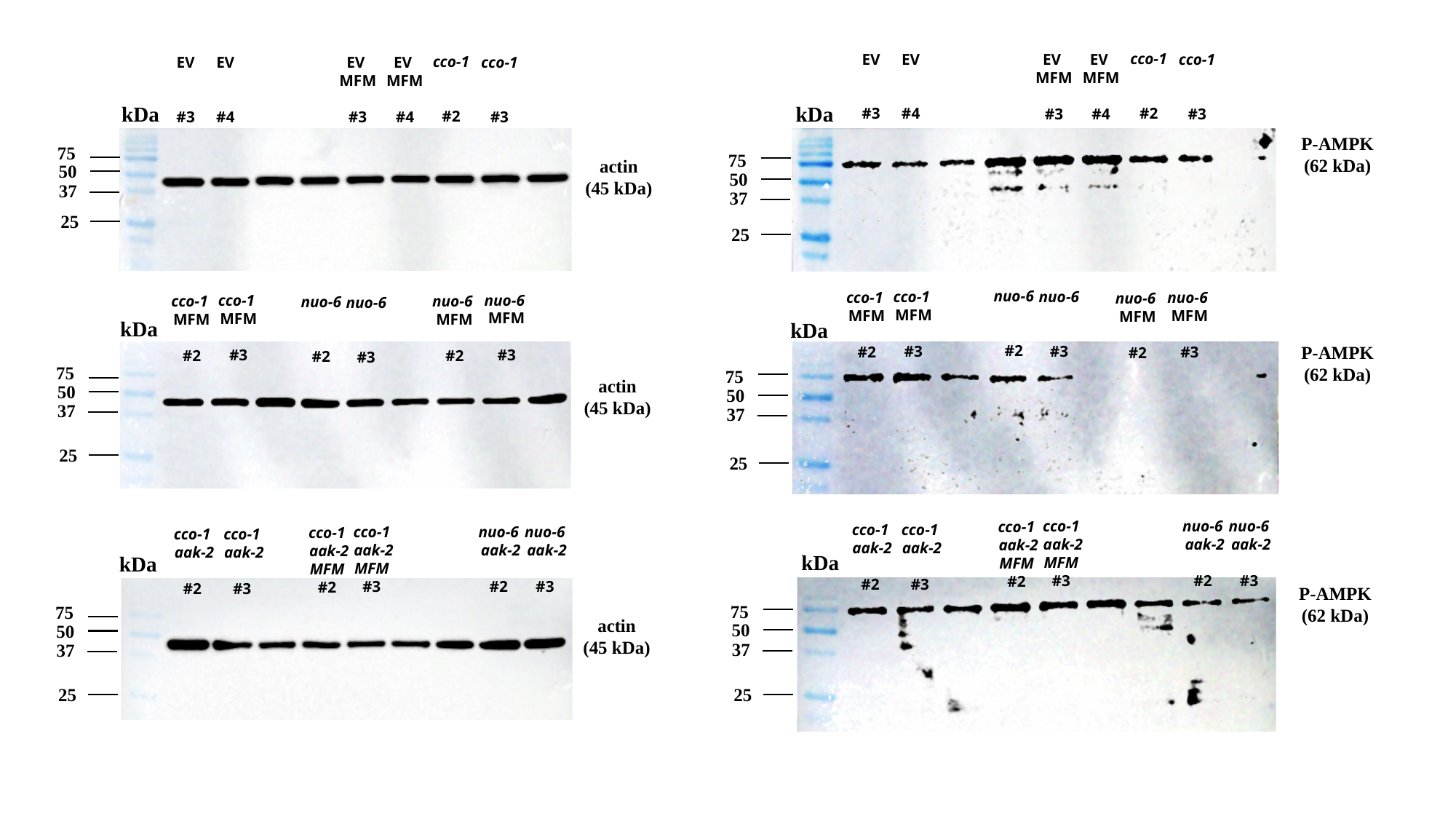

cco-1
#2
EV
#3
EV
#4
cco-1
#3
EV
MFM
#3
EV
MFM
#4
cco-1
#2
EV
#3
EV
#4
EV
MFM
#3
EV
MFM
#4
cco-1
#3
kDa
kDa
P-AMPK
(62 kDa)
75
75
50
50
37
37
25
25
nuo-6
#2
cco-1
MFM
#3
nuo-6
#3
nuo-6
MFM
#3
cco-1
MFM
#2
nuo-6
MFM
#2
nuo-6
MFM
#3
cco-1
MFM
#3
cco-1
MFM
#2
nuo-6
MFM
#2
nuo-6
#2
nuo-6
#3
kDa
kDa
P-AMPK
(62 kDa)
75
75
50
50
37
37
25
25
cco-1
 aak-2
MFM
#3
nuo-6
 aak-2
#2
nuo-6
 aak-2
#3
cco-1
 aak-2
MFM
#2
cco-1
 aak-2
#3
cco-1
 aak-2
#2
cco-1
 aak-2
MFM
#3
nuo-6
 aak-2
#2
nuo-6
 aak-2
#3
cco-1
 aak-2
MFM
#2
cco-1
 aak-2
#3
cco-1
 aak-2
#2
kDa
kDa
P-AMPK
(62 kDa)
75
75
50
50
37
37
25
25

### Slide 3
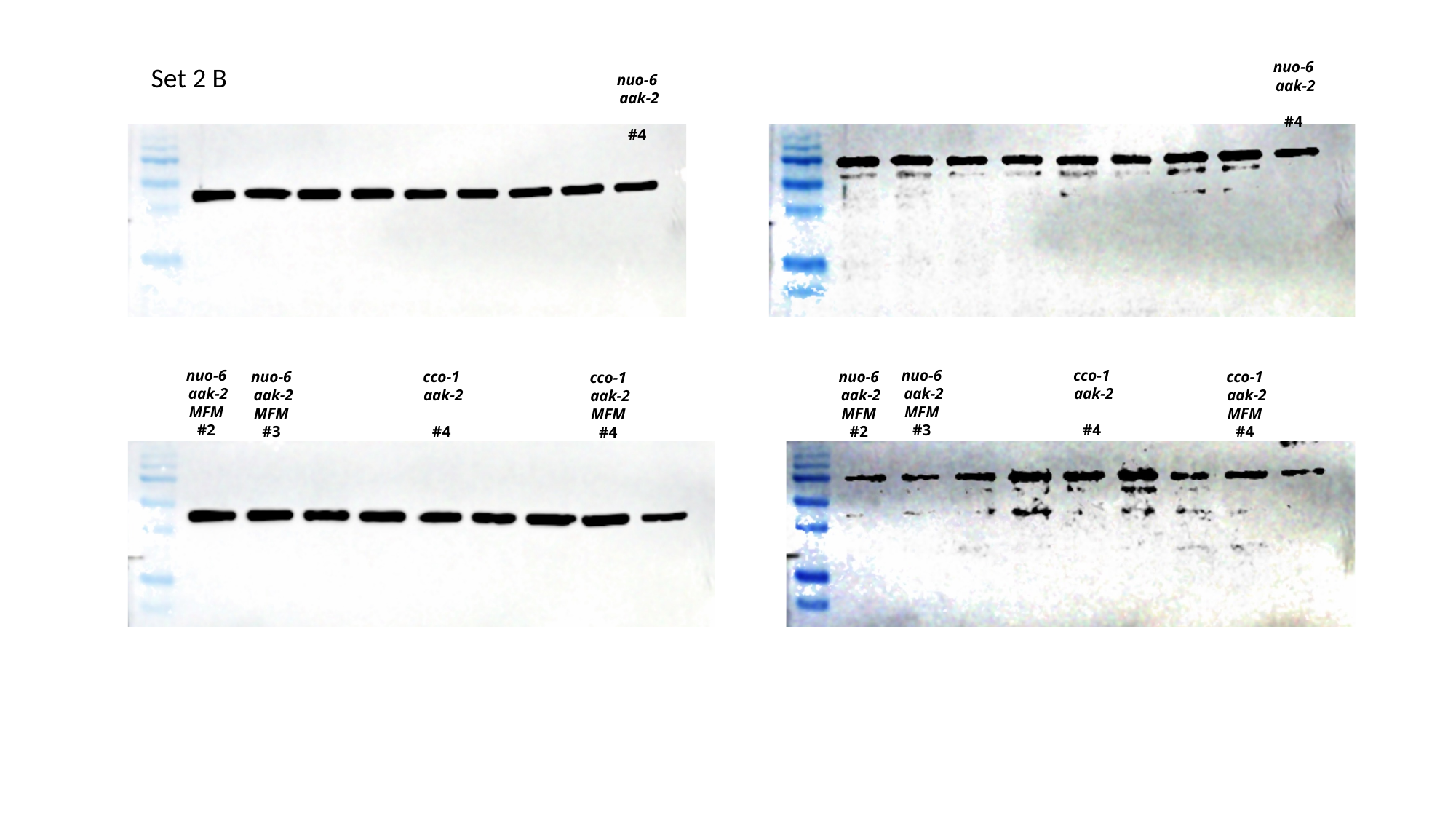

nuo-6
 aak-2
#4
Set 2 B
nuo-6
 aak-2
#4
nuo-6
 aak-2
MFM
#2
nuo-6
 aak-2
MFM
#3
cco-1
 aak-2
#4
nuo-6
 aak-2
MFM
#3
cco-1
 aak-2
#4
nuo-6
 aak-2
MFM
#2
cco-1
 aak-2
MFM
#4
cco-1
 aak-2
MFM
#4
